# Benchmarking substrate-based kinase activity inference using phosphoproteomic data

**DOI:** 10.1101/080978

**Authors:** Claudia Hernandez-Armenta, David Ochoa, Emanuel Gonçalves, Julio Saez-Rodriguez, Pedro Beltrao

## Abstract

**Motivation:** Phosphoproteomic experiments are increasingly used to study the changes in signalling occurring across different conditions. It has been proposed that changes in phosphorylation of kinase target sites can be used to infer when a kinase activity is under regulation. However, these approaches have not yet been benchmarked due to a lack of appropriate benchmarking strategies.

**Results:** We curated public phosphoproteomic experiments to identify a gold standard dataset containing a total of 184 kinase-condition pairs where regulation is expected to occur. A list of kinase substrates was compiled and used to estimate changes in kinase activities using the following methods: Z-test, Kolmogorov Smirnov test, Wilcoxon rank sum test, gene set enrichment analysis (GSEA), and a multiple linear regression model (MLR). We also tested weighted variants of the Z-test, and GSEA that include information on kinase sequence specificity as proxy for affinity. Finally, we tested how the number of known substrates and the type of evidence *(in vivo, in vitro* or *in silico)* supporting these influence the predictions.

**Conclusions:** Most models performed well with the Z-test and the GSEA performing best as determined by the area under the ROc curve (Mean AUC=0.722). Weighting kinase targets by the kinase target sequence preference improves the results only marginally. However, the number of known substrates and the evidence supporting the interactions has a strong effect on the predictions.

## Introduction

The functional plasticity of the proteome is modulated by specific post-translational modifications (PTMs) such as phosphorylation, glycosylation or ubiquitination. Reversible modification of proteins controls signal transduction and information processing within the cell, mediating molecular processes such as protein enzymatic regulation, complex associations, protein degradation and changes in subcellular localization (Choudhary & Mann 2010). Through the coordinated regulation of multiple parallel events, the cell is able to integrate all available signals to appropriately respond to environmental stimuli (Francavilla et al. 2016). Reversible phosphorylation of individual residues remains one the most studied PTMs and its status results from the net activity of kinases and phosphatases. Kinases bind covalently a phosphate group onto specific amino-acids, most frequently serine, threonine or tyrosine, while phosphatases catalyse the reverse reaction.

Previous studies of cell decision making mediated by protein phosphorylation have been limited by the reduced number of phosphosites and kinase activities that were possible to measure, most frequently using phospho-specific antibodies (Miller-Jensen et al. 2007). However, recent developments in large-scale phosphoproteomics and mass spectrometry (MS) have fostered the system-wide identification and quantification of phosphorylated sites before and after controlled perturbation (Olsen et al. 2006). These advances have not only allowed for the measurement of the response of thousands of individual phosphosites in a very rapid and granular time-frame (Kanshin et al. 2015; Humphrey et al. 2015), but also to estimate large scale enzymatic regulation based on the changes in known kinase substrates (Drake et al. 2012; Casado et al. 2013). The emergence of methods to estimate the activities directly from high-throughput MS data and prior knowledge in kinase substrate interactions introduces a new perspective on understanding the signaling response. These have been used to study differences in tumours (Drake et al. 2012; Casado et al. 2013), to study the effects of drugs, to reconstruct signaling networks and to broadly survey the kinase signaling states of the cell (Ochoa et al. 2016).

Substrate-based kinase activity prediction methods are founded on the assumption that the regulatory state of a kinase is reflected on the change of phosphorylation levels of its substrate sites. Drake and colleagues first reported this notion and performed an enrichment analysis of kinase activities (Drake et al. 2012). The method used consisted of a permutation analysis to infer an enrichment score using the Kolmogorov Smirnov statistic, an algorithm similar to the popular Gene Set Enrichment Analysis (Drake et al. 2012; Subramanian et al. 2005). This approach has been applied to a large number of kinases and conditions in a recent study (Ochoa et al. 2016). Alternatively, Casado et al. proposed an algorithm based on a one sample Z-test to measure kinase regulation in label-free cancer samples (Casado et al. 2013). More recently, the IKAP machine learning method models the abundance of each phosphosite as the sum of all kinase effects acting on it (Mischnik et al. 2016).

These methods depend on the aggregation of kinase substrate annotations. Comprehensive resources have aggregated experimentally verified *in vivo* and *in vitro* interactions between kinases and phosphorylation sites, including PhosphoSitePlus (Hornbeck et al. 2015), Phospho.ELM (Dinkel et al. 2011), HPRD (Peri et al. 2004) and Signor (Perfetto et al. 2016). This information is still limited to a fraction of all sites and kinases in very few model organisms. Computational approaches such as NetworKIN attempt to complete our knowledge on kinase targets by predicting new substrates from motif analysis and functional context derived from STRING (Kersten et al. 2009; Mering 2003).

The continuous development of new strategies requires a standardized framework to assess the substrate-based prediction of kinase activities. Therefore, we collected here a set of 24 conditional phosphoproteomic studies, in which 30 different kinases are expected to display regulation. This gold standard was used to evaluate 5 different methods: Z-test, Kolmogorov-Smirnov test, Wilcoxon ranksum test, gene set enrichment analysis algorithm (GSEA) and a multiple linear regression model (MLR). We also evaluated the effect of the kinase sequence specificity by weighting differently the target sites showing different sequence similarities to the binding motif. Moreover, we benchmarked the impact of the number of known target sites, as well as the effect of the evidence supporting the substrate interactions on the performance of the methods.

Our proposed gold standard and analysis provides a comprehensive comparison of the strengths and limitations of different methodologies and provides the necessary framework to evaluate future developments.

## Methods

### Defining a benchmark dataset for kinase regulation predictions

To evaluate the performance of different kinase activity prediction methods, we screened the literature to find 24 publications describing quantitative phosphoproteomic data reporting the response of a variable number of human phosphosites after perturbations linked to specific kinase activations or inhibitions. For example, the EGFR tyrosine kinase-receptor is activated when its ligand, the epidermal growth factor (EGF), binds the receptor (Oda et al. 2005). It is therefore expected that experiments assaying the phosphoproteomic response after EGF stimulation present increased activities of EGFR. Similarly, the molecular response to DNA damage has been associated with up-regulation of the ATR and ATM kinases (Matsuoka et al. 2007). Alternatively, drug treatments such as the addition of the kinase inhibitors Erlotinib or Torin 1 are expected to specifically inhibit their corresponding targets EGFR and mTOR. As a result of the curation effort, we collected a gold standard positive set containing 184 pairs between 30 different kinases regulated in 62 different perturbations with publicly available phosphoproteomic data (Table 1). A more detailed description of the conditions in the gold standard can be found in Table S1.

**Table 1.** Human kinases displaying specific regulation-up or down-in perturbations quantified using high-throughput phosphoproteomics.

| Experimental Condition | Kinases | Regulation | References |
| --- | --- | --- | --- |
| AKTi (Akt inhibitor VIII) ,AKTi (MK-2206),PI3Ki (GDC-0941), PI3Ki (PI-103),PIK3CA activation (H1047R) + inhibitor | AKT1 | down | (Wilkes et al. 2015; Wu et al. 2014) |
| PIK3CA activation (E545K),PIK3CA activation (H1047R), RG7356 | AKT1 | up | (Wu et al. 2014; Weigand et al. 2012) |
| Anti-CD3, anti-CD3 + anti-CD28, Differentiation (PMA) | PRKCA | up | (Nguyen et al. 2016; Salek et al. 2013; Rigbolt et al. 2011) |
| PKCi (BIM-1),PKCi (Gö-6976) | PRKCA | down | (Wilkes et al. 2015) |
| ATR inhibitor (VE-821) | ATR | down | (Šalovská et al. 2014) |
| CAMK2i (KN-62),CAMK2i (KN-93) | CAMK2A | down | (Wilkes et al. 2015) |
| Crizotinib | ALK | down | (Oppermann et al. 2013) |
| Dasatinib (50 nM) | MAPK3,ABL,LCK | down | (Pan et al. 2009) |
| DNA damage (Etoposide), DNA damage (Ionizing radiation), Early S (Thymidine), G1_S (Thymidine), Late S (Thymidine) | ATR, ATM | up | (Beli et al. 2012; Olsen et al. 2010; Dephoure et al. 2008) |
| EGF | AKT1, BRAF, EGFR, MAP2K1, MAP2K2, MAPK1, MAPK3, RAF1 | up | (Mertins et al. 2012; Engholm-Keller et al. 2011; Pan et al. 2009) |
| EGFRi (PD-153035),EGFRi (PD-168393), Erlotinib,Gefitinib | EGFR | down | (Wilkes et al. 2015; Weber et al. 2012) |
| EGF + SB202190 | MAPK14 | down | (Pan et al. 2009) |
| EGF + U0126, MEKi (GSK-1120212),MEKi (U0126),Selumetinib (AZD6244) | MAP2K1, MAP2K2 | down | (Wilkes et al. 2015; Stuart et al. 2015; Pan et al. 2009) |
| ERKi (ERK inhibitor),ERKi (ERK inhibitor II),Dasatinib (50 nM) | MAPK1 | down | (Wilkes et al. 2015; Pan et al. 2009) |
| Iloprost (5 nM) | PRKACA | up | (Beck et al. 2014) |
| Mitosis (AZD1152), Mitosis (MLN8054) | AURKA | down | (Kettenbach et al. 2011) |
| Mitosis (AZD1152) | AURKB | down | (Kettenbach et al. 2011) |
| Mitosis (BI2536),Mitosis (short Taxol + BI2536),BI 4834 (on Mitosis Nocodazole), PLK1 Inhibitor (G2r - BI2536 ) | PLK1 | down | (Halim et al. 2013; Grosstessner-Hain et al. 2011; Kettenbach et al. 2011) |
| mTORi (KU-0063794), mTORi (Torin-1),Rapamycin,Starved,Torin1 | MTOR | down | (Wilkes et al. 2015; Hsu et al. 2011) |
| PIK3CA activation (H1047R), PIK3CA activation (E545K) | PIK3CA | up | (Wu et al. 2014) |
| P70S6K1i (DG2), P70S6K1i (PF-4708671) | RPS6KB1 | down | (Wilkes et al. 2015) |
| ROCKi (H-1152), ROCKi (Y-27632) | ROCK1 | down | (Wilkes et al. 2015) |
| Sorafenib | PDGFRB, PDGFRA, BRAF, RAF1, FLT1, FLT3, FLT4 | down | (Zhuang et al. 2013) |
| Quizartinib | PDGFRA, FLT3 | down | (Schaab et al. 2014) |
| LRRK2-IN-1 (Endogenous LRRK2),LRRK2-IN-1 (LRRK2 overexpression) | LRKK2 | down | (Schaab et al. 2014; Luerman et al. 2014) |
| VEGF | FLT4, FLT1 | up | (Zhuang et al. 2013) |
| Vemurafenib (PLX4032) | BRAF | down | (Stuart et al. 2015) |

### Human quantitative phosphoproteomic data preprocessing and normalization

The human quantitative MS data used in this study is a subset of a previous compilation of human perturbation studies (Ochoa et al. 2016). Despite the heterogeneity of the biological samples, as well as the technical variation introduced by different experimental setups, high reproducibility is expected in the quantitative response of the direct kinase regulations included in the gold standard. As in the previous study, we applied a set of filters and quality control steps: (a) we restricted the analysis to peptides mapped to the Ensembl canonical transcripts ignoring other modified isoforms; (b) the log2 fold changes of phosphopeptides containing the same set of modifications were considered equal entities, in spite of differing in sequence coverages; (c) the log2 fold changes of the same phosphopeptides in different replicates were averaged; (d) to increase the interpretation of the fold changes as single PTMs, only monophosphorylated phosphopeptides were considered. Finally, after applying all filters, we excluded perturbations in the gold standard containing less than 1000 quantified phosphopeptides. Quantifications across conditions were quantile normalized to maximize reproducibility across samples. The quantification data for each phosphosite for the conditions in the gold standard can be obtained from Supplementary Table 2.

### Substrate-based kinase activity inference methods

For each perturbation in the gold standard, we calculated the changes in kinase activities based on the quantitative phosphoproteomic profile and the set of known kinase substrates. A total of 2818 manually curated interactions were compiled from PhosphoSitePlus (Hornbeck et al. 2015), Phospho.ELM (Dinkel et al. 2011) and Human Protein Reference Database (Peri et al. 2004). From this collection, 150 autoregulatory sites were excluded in order to prevent biases due to direct regulatory phosphorylations. The 5 predictors here explored can be classified according to their statistical properties as: parametric tests (Z-test), non-parametric tests (Kolmogorov-Smirnov test and Wilcoxon test), multiple linear regression models (MLR) and empirical computational approaches relying on data permutations (GSEA algorithm).

The parametric one-sample Z-test compares the mean fold change of all substrates of a given kinase to the mean and variance of all fold changes in the same condition. The non-parametric methods instead, assess the phosphorylation differences between the substrate and the non-substrate fold-change distributions using different metrics: the KS-test estimates the maximum distance between the cumulative probability distributions and the Wilcoxon test evaluates the rank differences among both distributions. The Z-test was implemented in R as previously described in (Casado et al. 2013) and the non-parametric tests were calculated using the available R functions. The Gene Set Enrichment Analysis algorithm (GSEA) uses a modified weighted Kolmogorov-Smirnov test to look for the enrichment on substrates of each kinase within the top or bottom of the ranked list of phosphosites. To estimate the significance, the enrichment score was compared against an empirical distribution of scores derived from 10000 random sets of substrates of the same size. To run the method, we used the in-house developed and freely available *ksea* R package (https://github.com/evocellnet/ksea).

Each of the aforementioned methods produces a p-value that summarizes the significance of the observed kinase regulation. In order to get a kinase activity score that indicates whether the kinase is increasing or decreasing in activity, we calculated the -log10 of the p-values and signed them based on the mean fold change of all substrate phosphosites. If in a given condition the target sites of a kinase are increasing in phosphorylation, the kinase activity is expected to be also increasing and vice-versa.

In addition to the tests above we predicted kinase regulation using a linear regression approach:

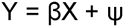

where, the dependent variable Y represents the phosphosite measurements in a condition and X is the connectivity matrix representing the associations with kinases. Xij equals 1 when phosphosite i is a known substrate of the kinase j, 0 otherwise. 4J represents the normally distributed error of the fit and are the weights of the model features and also represent the scores of the kinases activity changes in the condition. Kinase activities were estimated by solving the regression model applying a L2-norm regularization, ridge regression, and using the Python machine learning module scikit-learn (Pedregosa et al. 2011). Using a L2-norm regularization allows to ameliorate the impact of collinearity between features, namely different kinases can target similar sets of phosphosites. A regularization coefficient of 0.1 was used.

### Performance evaluation of the methods

To evaluate the performance of the kinase activity scores, we created a set of kinase-condition pairs containing all predicted pairs in the gold standard, as well as the same number of negative pairs generated by randomly sampling kinases and perturbations from the positive set. We repeated this procedure 60 times to generate a diverse space of negative interactions. Due to the incomplete coverage of the MS detection, only substrates for 24 kinases in the gold standard were quantified. Using the ranked lists of positives and the randomly generated negatives, the methods were evaluated using Receiver Operating Characteristic (ROC) and Precision-Recall (PR) curves. Area Under ROC Curve (AUC) and the observed precision at recall 0.5 were systematically used as performance metrics. The benchmark robustness was evaluated by comparing the summary metrics obtained in each of the 60 negative set randomizations. Additionally, the technical variance introduced by each of the methods was assessed by creating 10 permuted sets of kinase-substrate relationships maintaining the same number of substrates per kinase.

### Weighting kinase target sites by the kinase sequence specificity

The weighted Z-test and GSEA methods introduced in this study modify the original statistics to re-rank the substrate quantifications in order to include binding specificities as a proxy for their binding affinities. To calculate the estimated affinity, we constructed a library of position weight matrices (PWMs) from the amino-acid sequences surrounding the known kinase substrates (+/-7 residues). Next, we calculated the matrix similarity scores (MSS) to measure the identity between each substrate flanking sequence and their corresponding kinase-PWM (Wagih et al. 2015). The MSS were calculated using a re-implementation of the MATCH™ tool (Kel et al. 2003) adapted to consider aminoacid sequences (https://github.com/omarwagih/matchtm). The MSS-ranging from 0 to 1 - can be interpreted as an approximation of the kinase-substrate binding preferences. The kinase substrate fold changes were then multiplied by their corresponding MSS to obtain their weighted fold changes. As a consequence, differentially regulated substrates not showing affinity for the known kinase binding motif would see their fold changes diminished. The weighted fold changes were then used to perform the Z-test and the GSEA algorithm as described previously. The fold changes of the sites whose phosphorylation is not catalyzed by the kinase under prediction remained unaltered. When benchmarking the weighted version of the methods we only considered kinases with at least 10 known substrates available to build the PWM. This decreases the benchmark set to 135 positive kinase-condition pairs including 21 kinases covering 57 different perturbations.

### Benchmarking *in vivo, in vitro* and *in silico* substrates

Kinase activity prediction performances were evaluated separately depending on the support of the kinase-substrate relationships: *in vivo, in vitro* or *in silico.* The *in vivo* and *in vitro* sets were collected from the curated information available in PhosphositePlus (Hornbeck et al. 2015), while the *in silico* predictions were derived from NetworKIN 3.0 (Linding et al. 2007). NetworKIN integrates consensus substrate motifs based in NetPhorest (Miller et al. 2008) with functional contextual information derived from STRING (Mering 2003), in order to improve the kinase-substrate inference. A minimum of 3 substrates per kinase were required for the 3 sets of evidences. The negative sets in each case were generated using the same criteria as in the full set of substrates. To prevent the biases derived from the different number of available substrates in each category, the analysis was repeated by downsampling the substrates to the size of the smallest set 25 times.

## Results

### Defining a benchmark dataset of expected kinase regulation

The critical assessment of existing and forthcoming methods to infer changes in kinase regulation requires the creation of appropriate benchmark datasets. Based on prior knowledge in kinase regulation, we identified from a previous public compilation (Ochoa et al. 2016) a set of human quantitative phosphoproteomic datasets where the assayed perturbations are expected to trigger specific kinase responses. From all the assayed conditions, we defined a benchmark set of 184 positive kinase-condition pairs including 30 kinases and 62 experimental conditions. The selected perturbations include direct molecular regulation via kinase inhibitors or induction of cellular pathways as consequence of cellular processes or differentiation. The curated gold standard set of kinases regulated under MS-quantified conditions is shown in Table 1 along with the expected regulatory effect - up or downregulation. The total number of quantified phosphopeptides and more detailed information about the biological perturbations is available in Table S1. Although the gold standard list of expected regulations is unlikely to be complete due to downstream and "off-target" effects, it contains an enriched set of high confidence conditional kinase regulations where direct mechanisms of action have been reported in the literature.

### Substrate-based inference of kinase activity changes

In order to benchmark the changes in kinase activities under different biological perturbations, we compared 5 different methodologies that integrate quantitative phosphoproteomics data and the network of known kinase substrates. All of the predictors tested have as a premise that known substrates of kinases under regulation display a significant change in phosphorylation over the background changes occurring in all phosphosites in the same condition. We evaluated the following methodologies: Z-test, Kolmogorov-Smirnov test, Wilcoxon rank-sum test, a multiple linear regression (MLR) model and the gene set enrichment analysis (GSEA) algorithm (described in Methods). For all methods except the MLR, the -log10 of the p-value was used as a proxy of the kinase regulatory response. These values were signed according to the mean fold change of the kinase substrate sites, in order to estimate the kinase regulatory effect-activation or inhibition. For the MLR method we used the beta values of the fitted model to quantify kinase regulation.

### Benchmarking kinase activity changes predicted by different methodologies

For each method, we compared the absolute activity changes for kinases expected to display conditional regulation according to the gold standard, against the activities of the kinases in random conditions (Fig. S1). For all methods, kinases under expected regulation display changes in activity significantly higher than random kinases (two-sample Wilcoxon-test, all *p*-values <= 1.333e-06). It is noteworthy that some kinases in the negative set also present significant regulation under specific conditions, potentially due to other regulatory events not included in the gold standard. These could include, for example, downstream activation/inhibition of kinases or off-target effects in the cases of drugs. All methods were benchmarked using ROC and precision/recall analysis (Fig. 1 and Fig. S2). The GSEA and the Z-test yield comparable results showing higher median AUCs (0.724 and 0.721 respectively) and median precision values at recall 0.5 (0.725 and 0.708) than other approaches. To elucidate the performance variance due to technical variability, we compared the AUCs and precisions derived from the predictions based on the known regulons with those obtained randomizing the network of kinase-substrates maintaining the same degree distribution (Fig. S3). As expected, the randomization of the list of kinase targets results in near random predictions.

To evaluate the robustness of the methods to the number of quantified substrates, we compared the aforementioned performances with those obtained only for kinases with less or equal than 5 quantified substrates and those with more than 5 (Fig. 1). Overall, activity predictions of kinases with fewer quantified substrates present lower performances. When comparing performances across methods, Wilcoxon, KS and MLR present significantly lower AUC performances than GSEA or Z-test (Wilcoxon-test, p-value < 2.2e-16) if the number of quantifications remains below 5 substrates. All methods present similar performances for kinases with more than 5 known substrates, with GSEA and Z-test displaying the best results in all cases.

**Figure 1.**
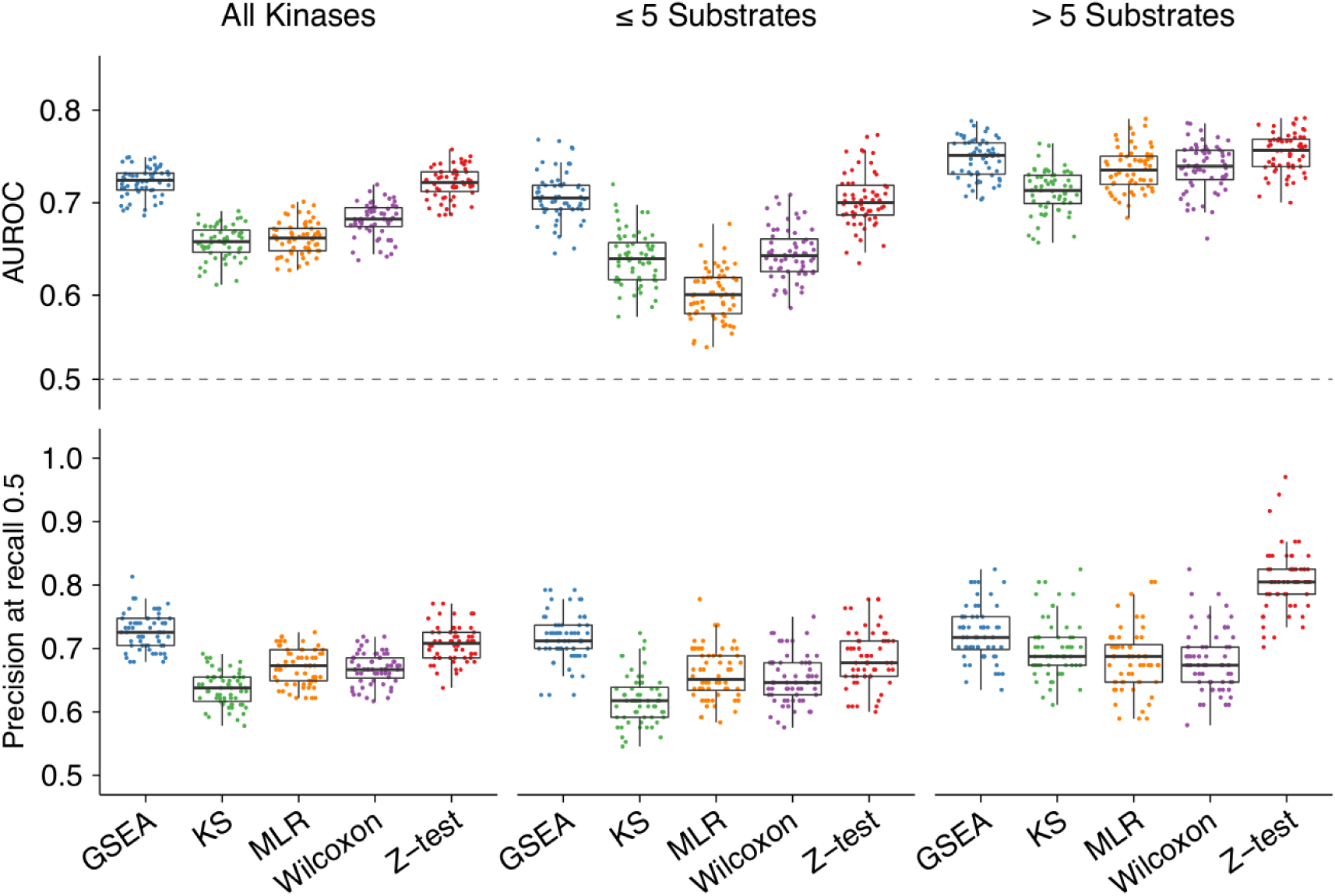
Comparison of kinase activity prediction performances by methodology and number of known kinase substrates. The filtered gold standard of 149 kinase-condition pairs was assessed against the same number of 60 randomly generated negatives (dots). Kinases are grouped by the number of quantified substrates: All, substrates <=5 (83 pairs) and substrates > 5 (66 pairs). Performance was evaluated using AUG values (upper panel) and precision at recall 0.504, 0.493 and 0.5 respectively (bottom panel).

### Kinase target site information impacts on the kinase regulation predictions

We reasoned that not all kinase substrates might be equivalent proxies of the upstream kinase activity. We hypothesize that target sites that best fit the known kinase sequence preferences could better represent the regulation of the catalytic kinase. To test this, we obtained position specific scoring matrices (PSSMs) that describe the binding specificity of the kinases in the positive set (Methods). These were used to score each target site based on their sequence and the values incorporated into the Z-test and GSEA (Methods). In these "weighted" versions of the methodologies, phosphosites that best match the kinase preference were more strongly considered when predicting the regulation of the kinase. We observed a small but significant improvement in AUC and Precision at recall 0.5 whereby these changes tended to perform better than the initial implementations: Z-test-weighted vs Z-test (Wilcoxon-test, p-value = 8.415x 10−^10^) and GSEA-weighted vs GSEA (Wilcoxon-test, p-value = 1.266x 10−^6^)(Fig 2 and Fig S4).

**Figure 2.**
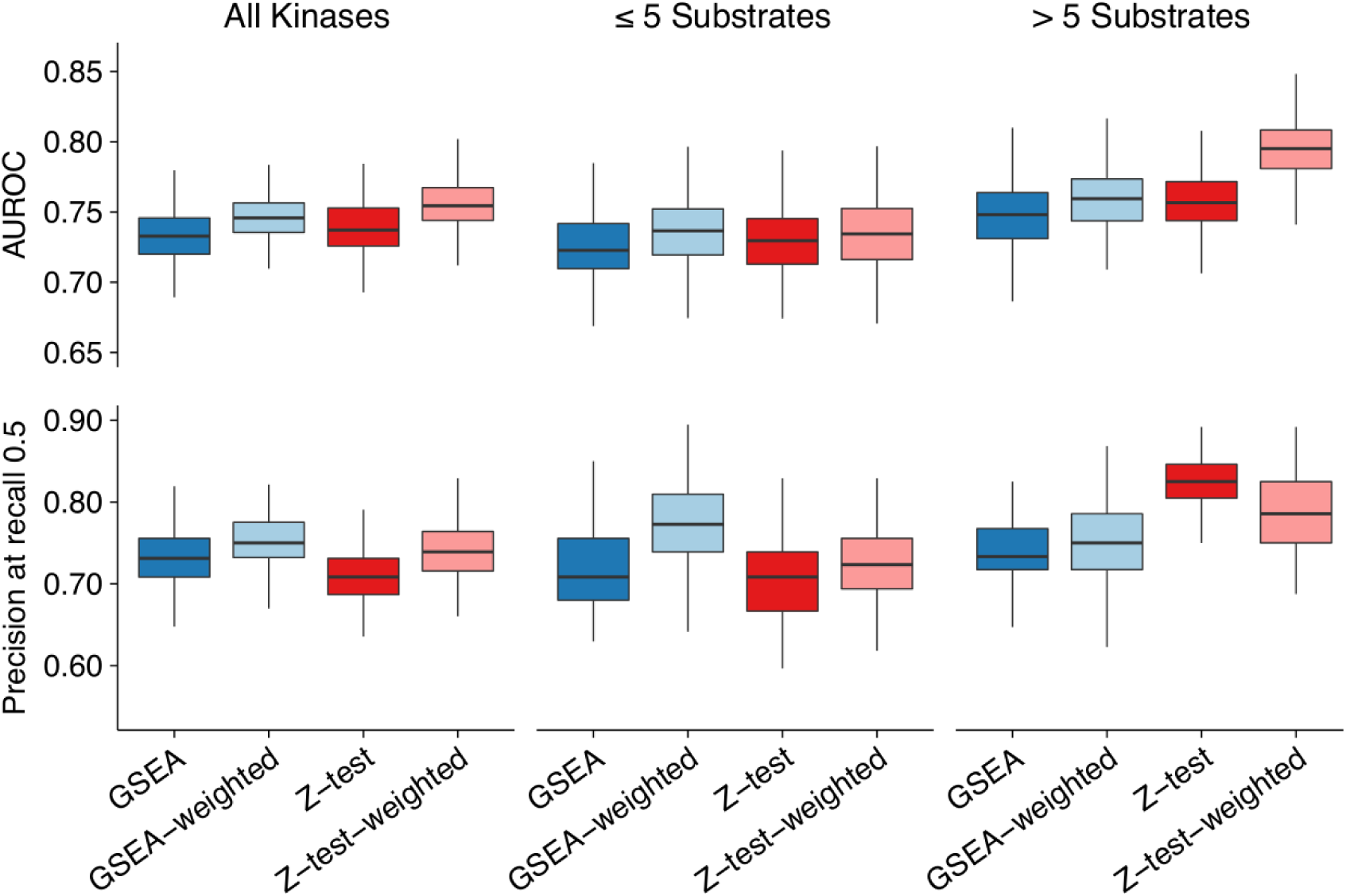
Comparison of substrate-based kinase activity prediction methods weighting substrates by kinase binding specificity. Similar to Figure 1. "Weighted" GSEA and Z-test include the similarity of the neighboring region to the kinase recognition motif to correct the relevance of each of the substrates (see Methods). Weighted and unweighted performances are based on the same set of kinases and conditions (135 positive kinase-condition pairs).

### Activities inferred using *in vivo, in vitro* or *in silico* supported substrates

In addition to analysing the effect of the substrate confidence based on their kinase specificity, we also compared the impact of the evidence supporting the kinase-substrate interactions. Candidate kinase substrates were separated into *in vivo, in vitro* and *in silico* supported substrates. Kinase activities predicted using GSEA or Z-test using *in vivo* characterized substrates perform better than predictions using *in vitro* supported substrates (AUC Wilcoxon-test, *p-value* < 2.2e-16) (Figure 3). Moreover, kinase activities based on substrates determined *in silico* display poor performances. To avoid potential biases derived from the different number of substrates in each set, we repeated the comparison by downsampling the substrates to the size of the smallest set (Fig S5). Also in this case, *in vivo* characterized substrates present better performances. Similarly, we restricted the comparison to those positive pairs predicted in all sets observing a similar trend (Fig S6). The poor performance displayed by the *in silico* substrates can be improved by selecting a more stringent prediction score (Fig S7). However, this effect is mostly due to an enrichment of previously characterized substrates, rather than the finding of novel substrates relevant for the kinase activity prediction. Our results highlight the relevance of the curated *in vivo* substrates to accurately infer changes in kinases activities.

**Figure 3.**
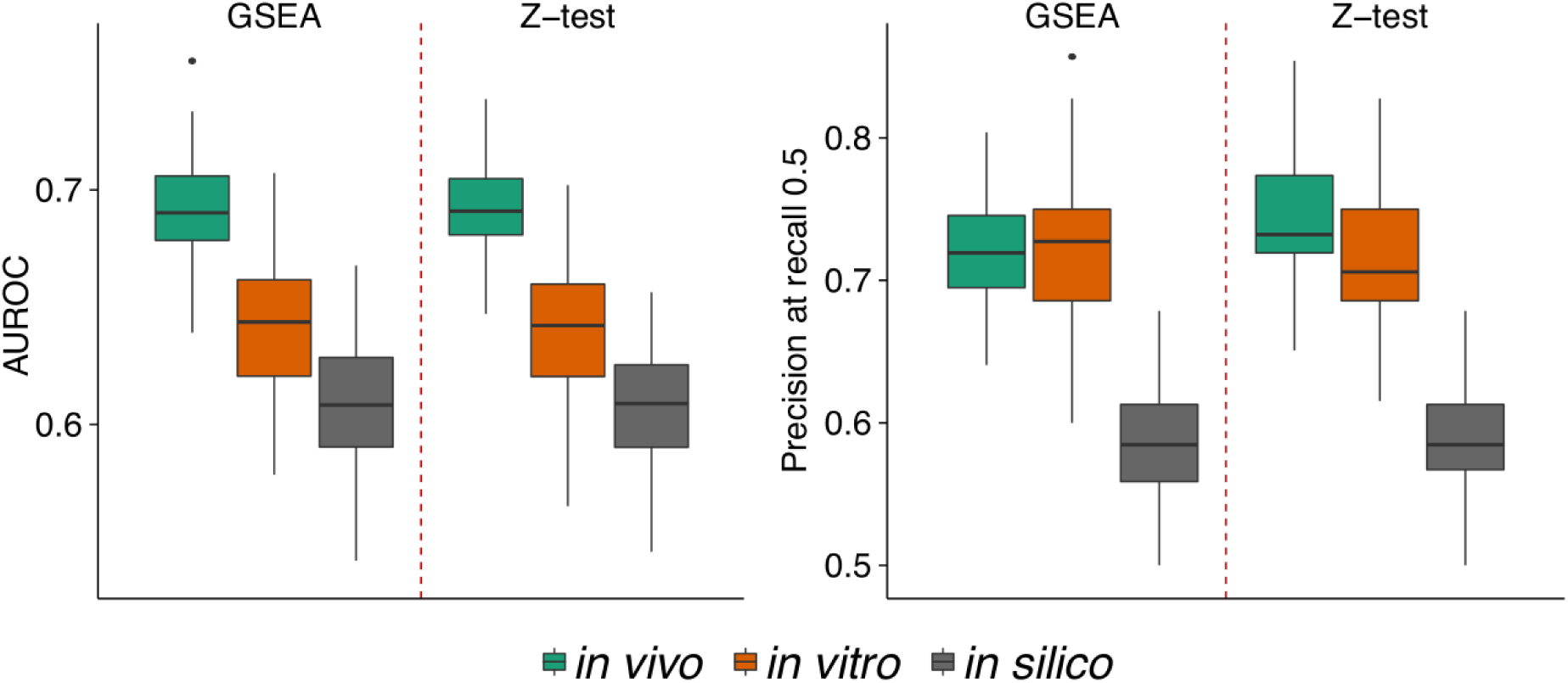
Benchmark of kinase activity predictions using *in vivo, in vitro or in silico* supported substrates. (A) AUC performance and (B) precision at recall 0.5. Positive kinase-condition pairs are 82, 48 and 75 respectively. The trend is maintained when downsampling the number of substrate sites to guarantee a similar number of target sites for all cases (Supplementary Figure S5) or restricting to the same set of kinases and conditions (Supplementary Figure S6).

## Discussion

The progress and developments in mass spectrometry have led to the unbiased exploration of cell signalling changes via phosphoproteomics. Single experiments are now reporting how changes of thousands of phosphorylation sites occur across tens to hundreds of conditions (Abelin et al. 2016; Mertins et al. 2016) and it has been proposed that the activation status of kinases can be inferred by the measurement of its targets sites (Drake et al. 2012; Casado et al. 2013). This approach has been used in several contexts, including the stratification of cancer patients (Drake et al. 2016), or to study kinase pathway changes occurring after specific stimulations (Terfve et al. 2015). Although it is intuitive, that change in abundance of target phosphosites reflects the activation status of kinases, such approaches have not yet been thoroughly benchmarked and compared. Here, we have collected a benchmark set of conditions where specific kinases are expected to be regulated and use it to compare different methodologies to predict kinase activity changes, as well as the quality of the list of substrates used. The required data is available as supplementary materials allowing others to easily compare alternative approaches. We also provide an R package including one of the best performing methods (GSEA at http://github.com/evocellnet/ksea).

Overall, we see a strong performance of all methods tested. Some true kinase-conditions pairs were not predicted to be regulated (Supplementary Figure 1). This could be due to technical issues in the treatments to stimulate and monitor the activity of the kinases, for instance: incorrect times selected to detect the changes in activity, inadequate stimulation of the control condition or defective concentration of the ligands to generate a cell response, etc. However, it should also be considered that complex regulatory mechanisms are involved in the control of kinase activation like feedback regulation, changes in subcellular localization or binding to scaffold proteins. It is possible that different sets of substrates (e.g. in different cellular compartments) might be relevant to estimate the activity for different conditions. This knowledge opens the door for the rational selection of kinase target sites that may serve as the best proxies for the activity of the effector kinase.

We observed dependence on the evidence supporting the kinase substrate interaction. This cautions against the use of predicted kinase interactions in such algorithms, at least with their current state of the art. Reversely, this suggests that the co-regulation patterns of phosphosites across conditions can be used as a signal for the prediction of new kinase targets.

## Acknowledgements

PB acknowledges support from an HFSP CDA award (CDA00069/2013)

## Tables and Figures

**Supplementary Figure S1.**
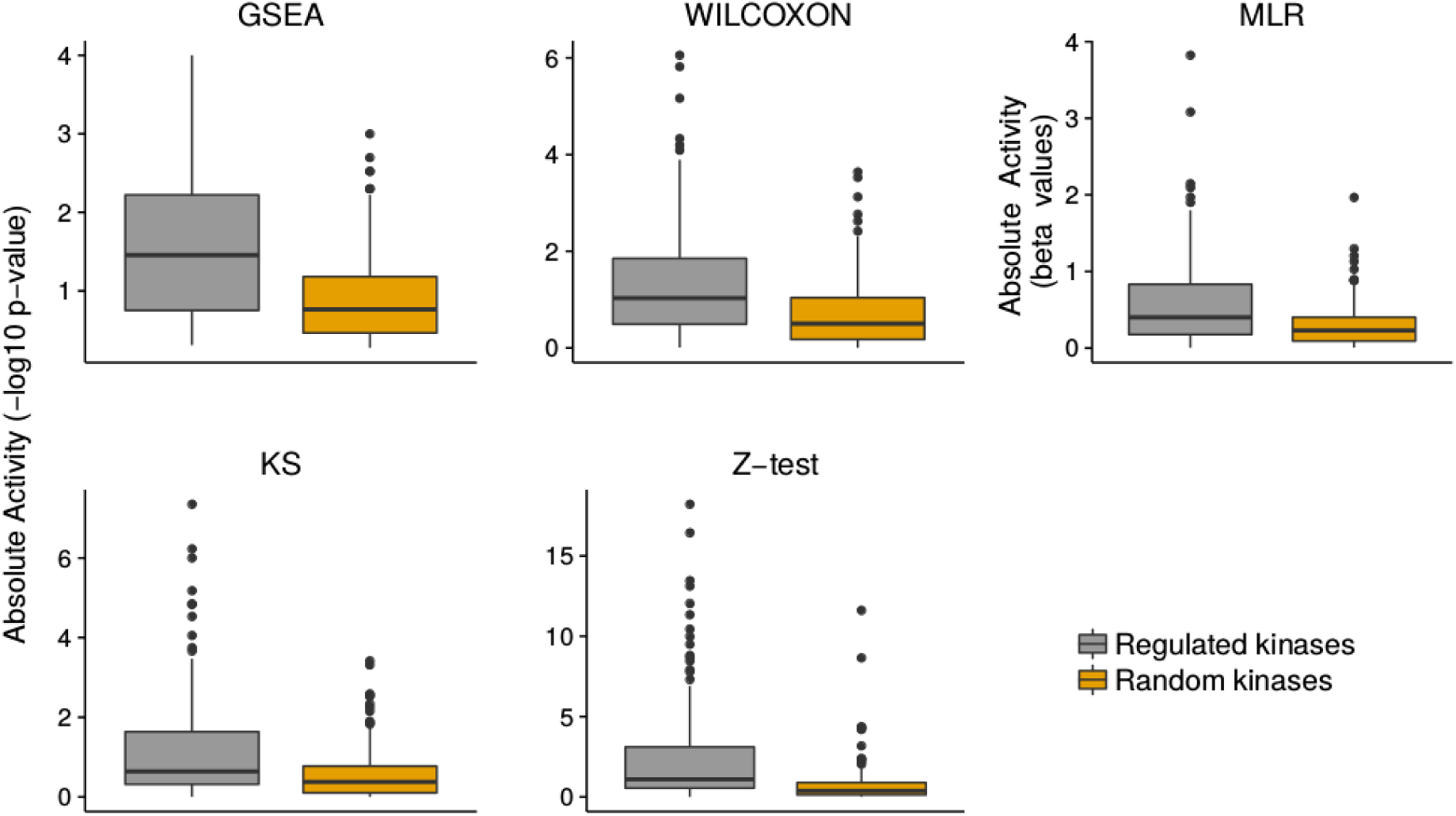
Predicting kinase regulation using 5 substrate-based methodologies. Absolute activity scores for 149 kinase-conditions expected to show regulation are compared against the same number of randomly generated pairs using the 5 methodologies under comparison.

**Supplementary Figure S2.**
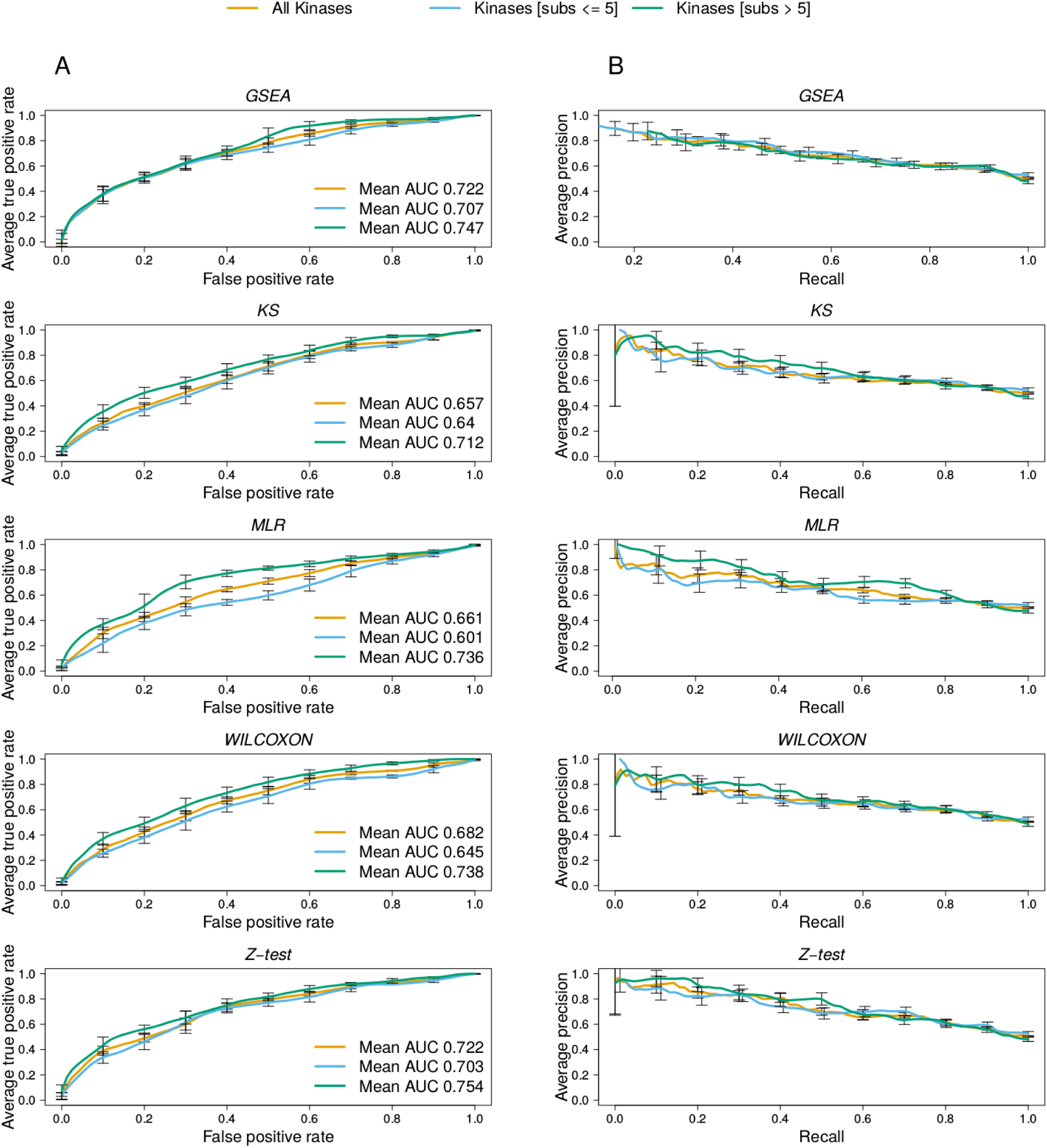
ROC and Precision-Recall (PR) curves comparing kinase activity prediction methods performance per number of known substrates. The kinase activity scores were calculated using the -log10 of the p-values of each statistic, with the exception of **MRL,** and were signed based on the mean phosphorylation value of the substrates set. For the **MRL** model the beta values were used as the estimation of the regulation of each kinase. A total of 149 kinase-conditions cases are present in the positive set and were paired with 60 negative sets of 149 independently sampled random kinase-conditions. (A) The ROC and **(B) PR** curves were calculated for the whole benchmark and by grouping the kinases in two categories: few(<= 5 substrates) and many substrates (> 5 substrates). The average curves of the 60 validation sets are shown for each comparison.

**Supplementary Figure S3.**
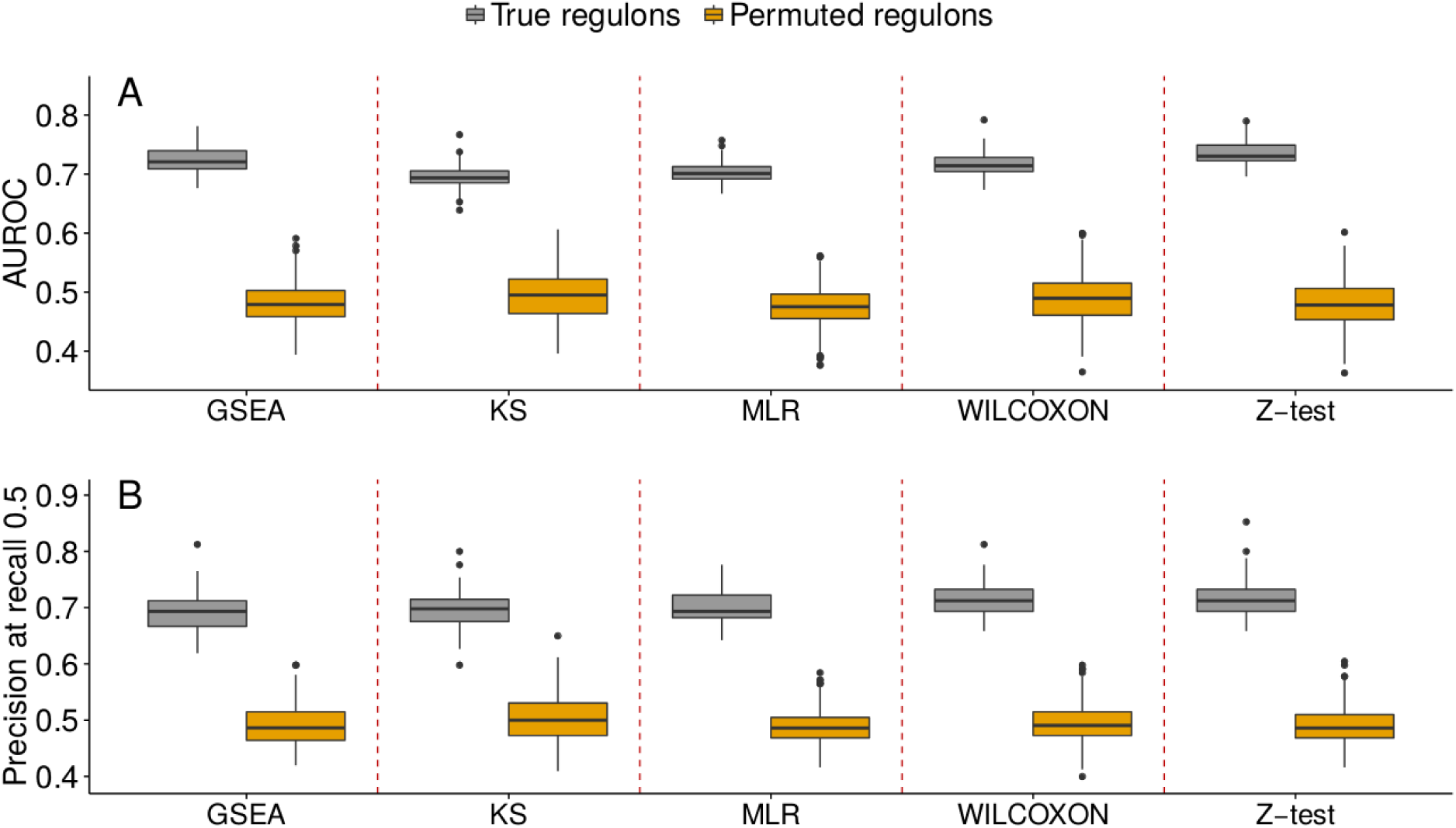
Methods performance comparison using the true and permuted sets of kinase-substrates interactions. A total of 10 permuted sets of kinase-substrate relationships conserving their original degrees of connectivity were generated to test the accuracy of the methods to random groups of substrates. The performance metrics of the methods were compared against the true kinase-substrates interactions. Only 105 positive cases of the benchmark described in Table 1 were considered due to missing quantifications of substrates in the permuted sets. This set was compared against 60 randomly generated negative sets to calculate the ROC and Precision-Recall curves. AUC values of the ROC curves (up) and precision values at recall 0.5 (down).

**Supplementary Figure S4.**
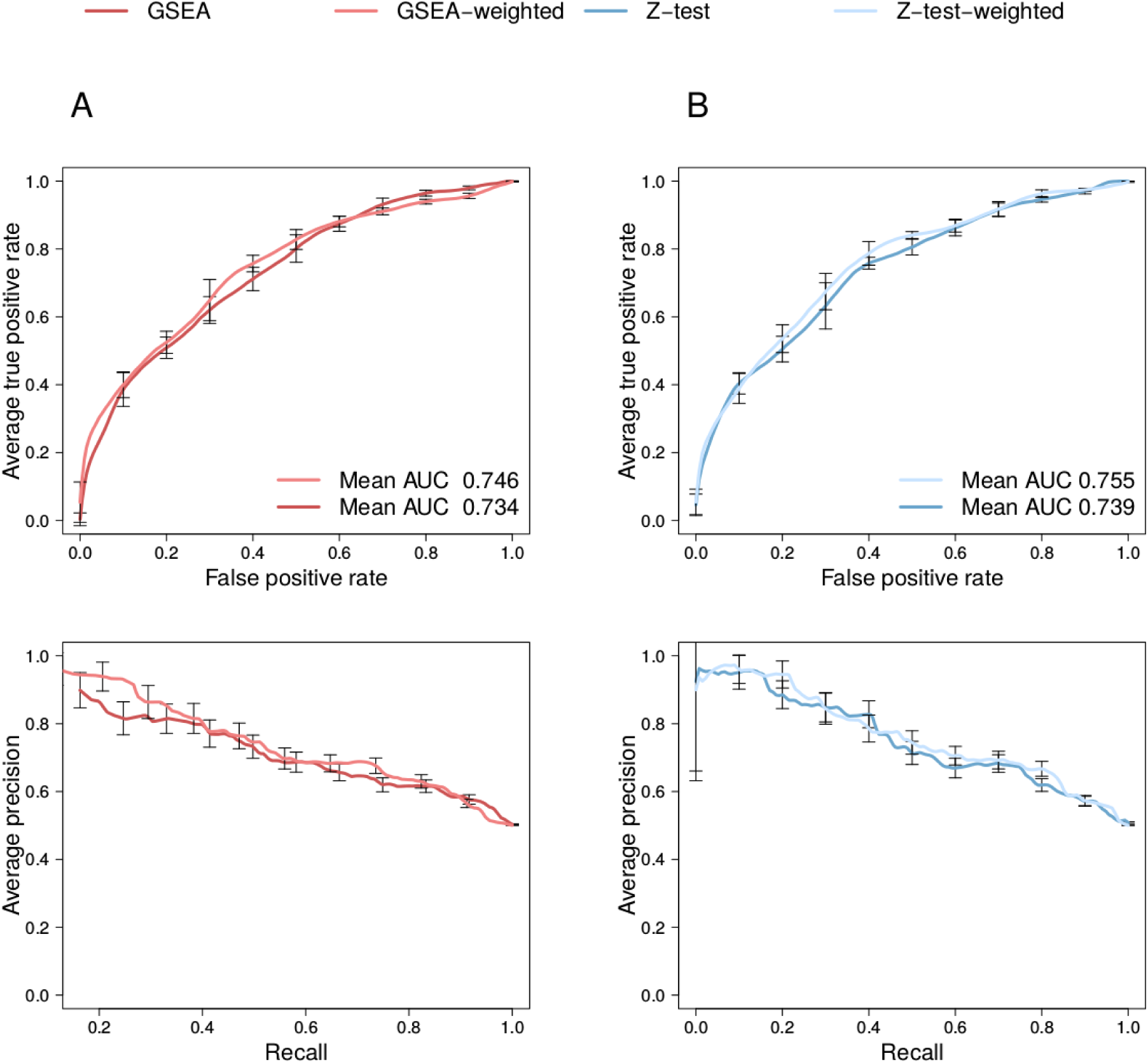
ROC and Precision-Recall (PR) curves for the weighted vs unweighted. Same data as Figure 2. (A) The ROC and (B) PR curves. The average curves of the 60 validation sets are shown for each method. A total of 135 kinase-conditions cases are present in the positive set and were paired with 60 random negative sets sampled independently.

**Supplementary Figure S5.**
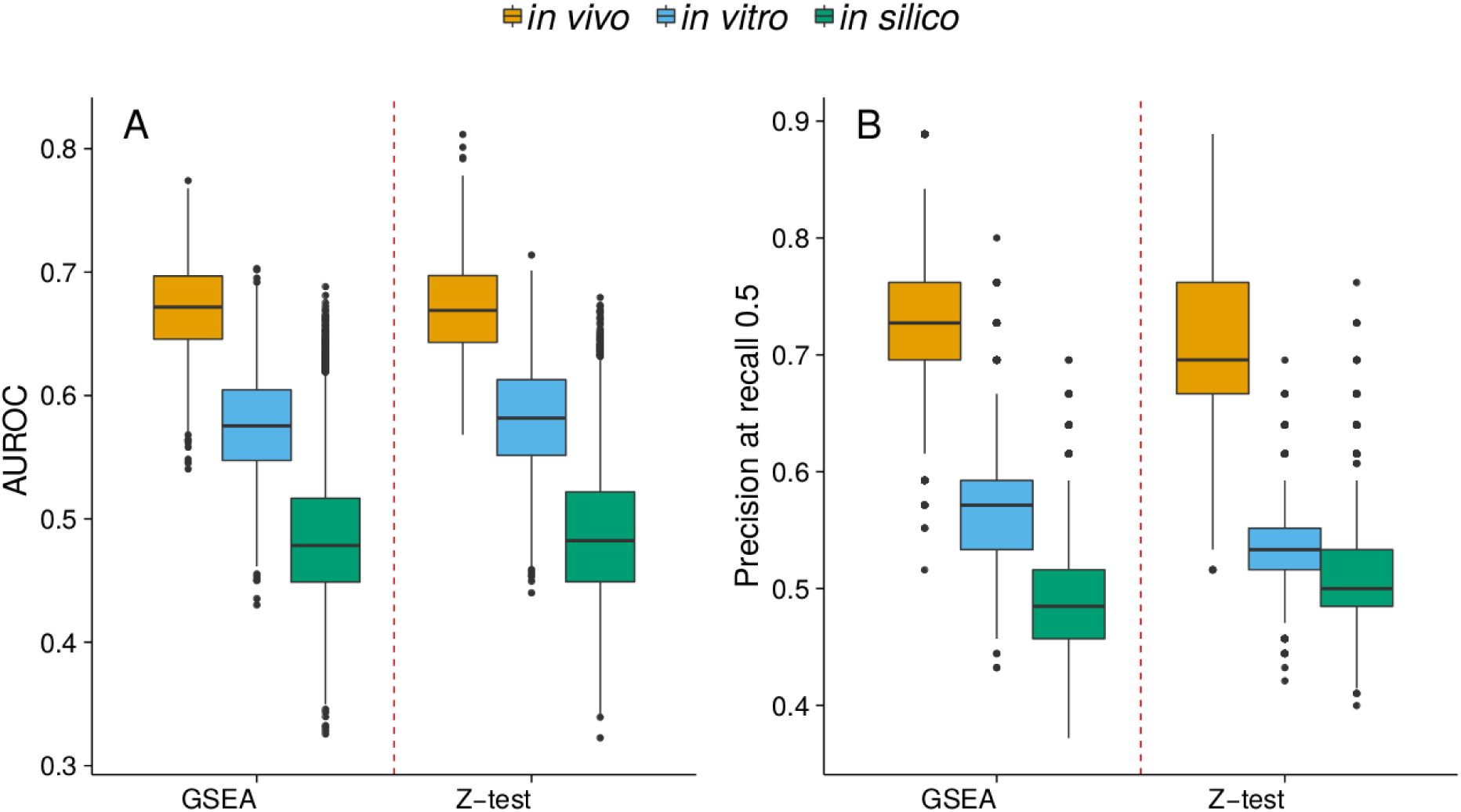
As Figure 3 but downsampling the substrates. Benchmark of kinase activity predictions using *in vivo, in vitro* or *in silico* supported substrates. (A) AUC performance. (B) Precision at recall 0.5. Positive kinase-condition pairs are 31 in all sets. Kinase-substrates collections were down sampled 25 times to the size of the smallest set.

**Supplementary Figure S6.**
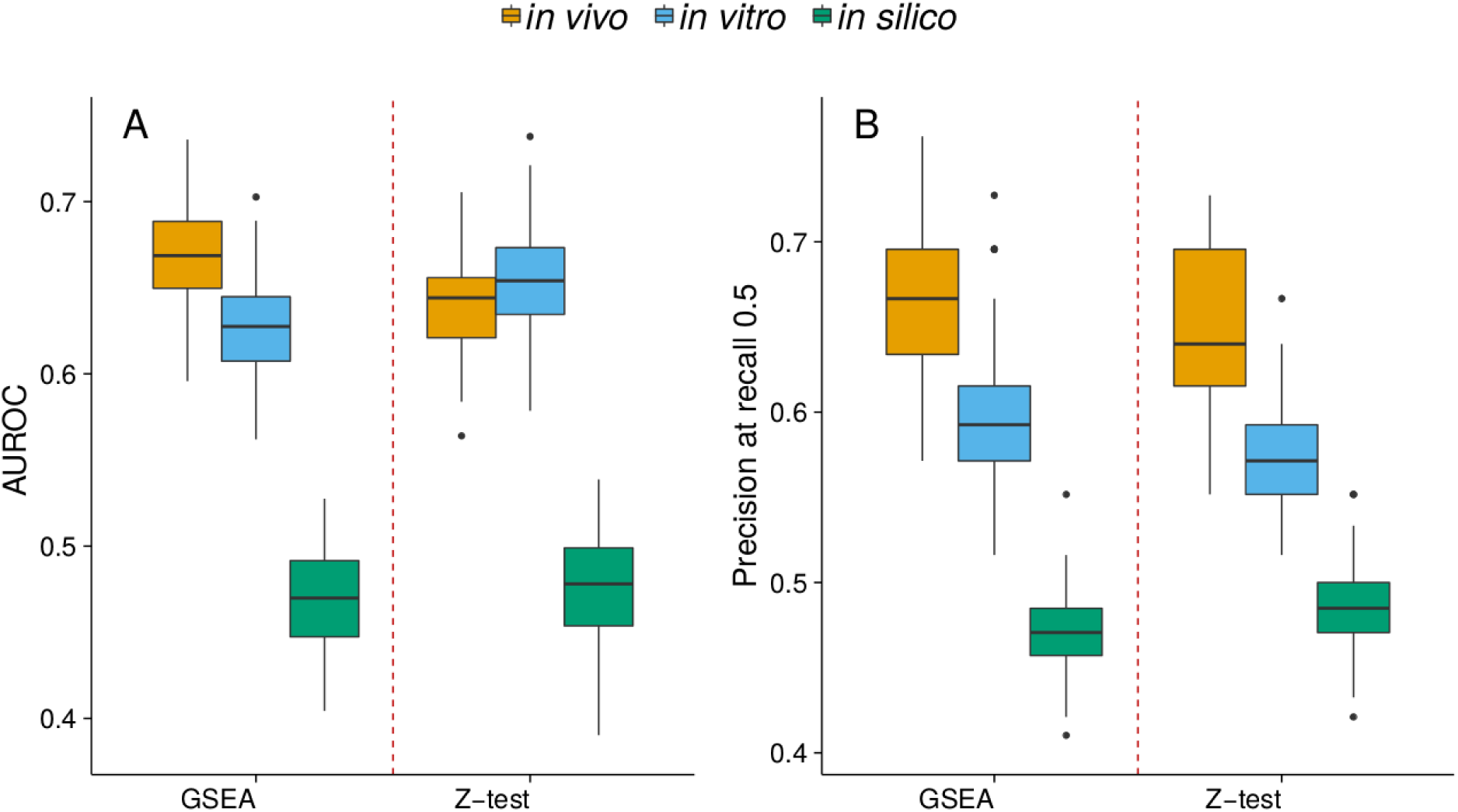
As Figure 3 but having the same positive in all sets. Benchmark of kinase activity predictions using *in vivo, in vitro* or *in silico* supported substrates. (A) AUC performance. (B) Precision at recall 0.5. Positive kinase-condition pairs are 31 in all sets.

**Supplementary Figure S7.**
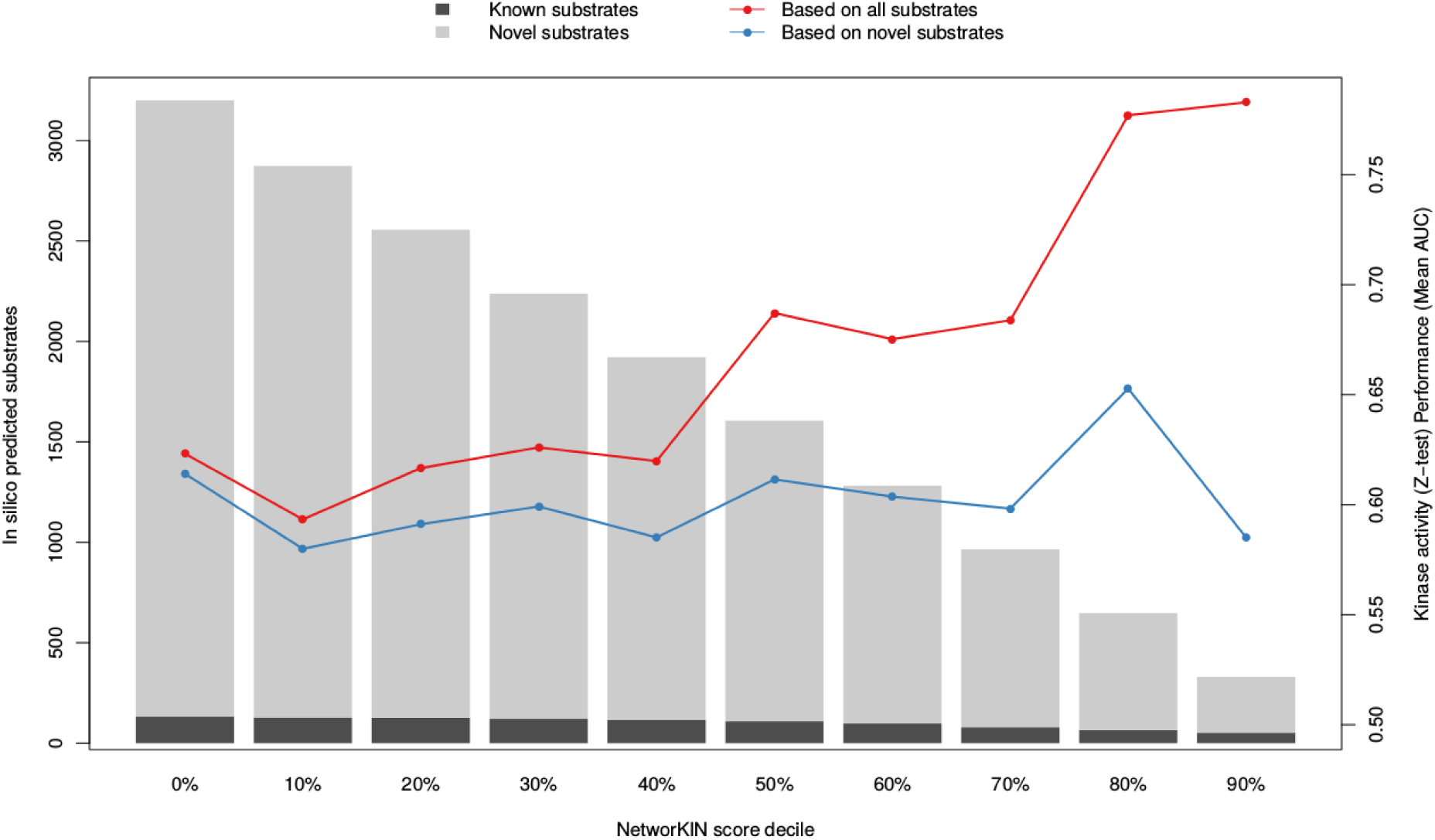
Effect of NetworKIN score in the performance on kinase activity predictions based on *in silico* supported substrates. NetworKIN predicted substrates are discretized per kinase using score deciles. Total number of known and previously reported NetworKIN predicted substrates are visualized in the barplot-left axis. Performance of the predictions measured as mean AUC of the Z-test predictions is displayed for all predicted substrates-red - and for all predicted substrates not previously reported-blue.

## Supplementary Tables

Table S1.Description of studies and biological perturbations included in gold standard.

Table S2.Quantifications of experimental perturbations included in gold standard.

